# Development of an enhanced anti-pan-N-formylmethionine-specific antibody

**DOI:** 10.1101/2024.12.12.628262

**Authors:** Dasom Kim, Kyu-Sang Park, Cheol-Sang Hwang

## Abstract

Both bacterial and eukaryotic ribosomes can initiate protein synthesis with formylmethionine (fMet), but detecting fMet-bearing peptides and fMet-bearing proteins has been challenging due to the lack of effective anti-pan-fMet antibodies. Previously, we developed a polyclonal anti-fMet antibody using a fMet-Gly-Ser-Gly-Cys pentapeptide that detects those fMet-bearing peptides and fMet-bearing proteins regardless of their sequence context. In this study, we significantly improved the antibody’s specificity and affinity by using a mixture of fMet-Xaa-Cys (fMXC) tripeptides (Xaa, any of the 20 amino acids) as the immunogen. This newly optimized anti-fMet antibody is a powerful, cost-effective tool for detecting fMet-bearing proteins across species. Furthermore, this approach provides a foundation for developing anti-pan-specific antibodies targeting other N-terminal modifications through acylation, alkylation, oxidation, or arginylation, etc.

**METHOD SUMMARY:** fMet-Gly-Ser-Gly-Cys (fMGSGC), fMet-dPEG4-Cys (fMdPEG4C), and fMet-Xaa-Cys (fMXC; Xaa, any of the 20 amino acids) were used as antigens to generate anti-pan-fMet-specific antibodies (anti-fMet antibodies). The quality of the raised antibodies was evaluated by immunoblotting using lysates from *Escherichia coli* (*E. coli*) DH5α cells and human kidney HK2 cells, as well as by enzyme-linked immunosorbent assay (ELISA) with purified fMet-bearing (fMet-) proteins and their unformylated counterparts.

## 1. Background

The N-terminus of proteins is highly reactive and exposed to various modifications, including proteolytic cleavage, acylation, alkylation, oxidation, and arginylation (1–7). These N-terminal (Nt-) residue-specific modifications often alter the overall protein charge, localization, stability, and interactions, ultimately affecting cellular stress response, protein aggregation, and cell fate (1,4,8–14). Unlike other co- and post-translational modifications, Nt-Met formylation occurs pre-translationally (before protein synthesis) in bacteria and organelles of bacterial origin, such as mitochondria and plastids (15–17). Specifically, formyltransferase (FMT) catalyzes the transfer of a formyl group from 10-formyltetrahydrofolate (10-THF) to Met linked to initiator tRNAi, producing formylated fMet-tRNAi. Then, fMet-tRNAi is incorporated into proteins during translation initiation, allowing nearly all newly synthesized proteins to retain Nt-fMet (15,16). However, as fMet-polypeptides emerge from the ribosome’s exit tunnel, ribosome-bound peptide deformylase (PDF) removes the Nt-formyl group, leaving an unmodified Met at the N-terminus (18–20). The counteracting function of PDFs makes the retention of fMet transient, significantly reducing the species and levels of fMet-proteins (20). Impaired deformylation, however, leads to toxic aggregation of fMet-proteins and prevents the essential cotranslational excision of the deformylated Nt-Met residue (21). As a result, fMet-proteins comprise only about 5% of mature bacterial proteins (1,22). In mitochondria and chloroplast, the population of fMet-proteins has not been fully explored, except for highly-formylated human COX1 (23,24).

In contrast to the long-lasting assumption that fMet-polypeptide production would be restricted to bacteria and bacterial-origin eukaryotic organelles (15), the recent studies have revealed that nuclear DNA-encoded fMet-polypeptides are also produced in the cytosol of yeast and human cells (2,25,26). Some of these fMet-proteins are shown to be selectively degraded by a eukaryotic fMet/N-degron pathway that recognizes Nt-fMet as a degradation signal (25). (Notably, although the role of Nt-fMet as a degradation signal has initially been proposed in bacteria, its corresponding destruction machinery has yet to be identified (17).) Furthermore, deformylation of cytosolic nascent fMet-polypeptides plays critical roles in yeast adaptation to undernutrition and cold stress (25), while also regulating colorectal cancer cell proliferation, suppressing cancer stem cell (CSC) properties, and influencing *in vivo* tumorigenesis (26).

Nt-Met formylation is involved in various biological processes beyond protein synthesis and degradation, such as complex formation, stress and immune responses, and human diseases. In human cells, mutations in mitochondrial formyltransferase (MTFMT), which reduce Nt-Met formylation, are linked to Leigh syndrome, a severe neurometabolic disorder (27), as well as with compromised mitochondrial integrity and increased susceptibility to intracellular infections (28). Elevated levels of fMet or fMet-peptides in human blood have been associated with late-onset diseases and increased mortality in septic shock and critical illness (29–31). Although these fMet derivatives are believed to originate from human microbes or mitochondria (29–31), cytosolic fMet-protein synthesis is likely the primary source, given the ability of cytosolic ribosomes to produce fMet-proteins and the large number of nuclear DNA-encoded proteins (2,25,32). Therefore, Nt-Met formylation greatly enhances the functional complexity of the cellular proteome (22) and plays crucial roles in various physiological and pathological processes (32).

However, the limited availability of tools beyond mass spectrometry for detecting a wide range of fMet-proteins and fMet-peptides has hindered the understanding of their functions and regulations across diverse organisms, from bacteria to humans (25,33). Although antibodies targeting specific fMet-peptides have been developed, their use has been restricted to proteins and peptides with sequences closely matching the antigen peptides employed in their production (25,34). The previous efforts to generate a pan-fMet-specific antibody using the pentapeptide antigen fMGSGC faced challenges in sensitivity and versatility (2). In this study, we addressed these limitations by designing a mixture of tripeptide antigen fMXC with all 20 amino acids at the second position, producing with a dramatic improvement in the performance of the newly developed anti-pan-fMet-specific antibodies.

## 2. Materials & methods

### 2.1. Generation of antisera

The Nt-formylated synthetic peptides fMGSGC, fMdPEG4C, and fMXC were, respectively, conjugated to the keyhole limpet hemocyanin (KLH) as the carrier protein. Sulfo-SMCC (M6035; Merck, NJ, USA) was used as the crosslinker between the peptides and KLH. Rabbit polyclonal antisera against each antigen were produced through a series of procedures: a primary injection at week 0, followed by boosting at weeks 4, 6, and 8, with final serum collection via heart puncture at week 9. The production of antisera was conducted by AbClon (Seoul, Republic of Korea).

### 2.2. Preparation of bacterial cell extracts

*Escherichia coli* DH5α was inoculated into an LB medium (25 g/l) and cultured overnight at 37°C. The following day, the culture was diluted 100-fold in fresh LB medium and grown until it reached A_600_ of 0.6. Actinonin (HY113952; MedChemExpress, NJ, USA), a PDF inhibitor, was dissolved in ethanol (20 mg/mLl) and added to the culture at a final concentration of 2 µg/ml for 30 min. In the mock group, sole ethanol without actinonin was treated instead. Following treatment, the culture was centrifuged at 12,000*g* for 2 min at 4°C to collect *E. coli* pellets. The pellets were lysed in a 2% sodium dodecyl sulfate (SDS) (BR1610302; Bio-Rad, CA, USA) solution containing a protease inhibitor cocktail (4693116001; Merck) and 20 µg/ml actinonin, then incubated on ice for 20 min, sonicated briefly, and centrifuged at 12,000*g* for 10 min at 4°C to obtain clear lysates. The protein concentration of the lysates was measured using a BCA protein assay kit (23250; Thermo Fisher Scientific, MA, USA). The lysates were diluted to a concentration of 2 µg/µl, mixed with a 4× SDS sample buffer in a ratio of 3:1 (v/v), and heated at 70°C for 10 min to prepare samples for Coomassie brilliant blue (CBB) staining and immunoblotting.

### 2.3. Preparation of human cell extracts

The human HK2 cell line, derived from a human kidney, was cultured using a DMEM/F-12 medium (11320033; Thermo Fisher Scientific) supplemented with 10% FBS (SH30919.03; Cytiva, MA, USA). Once the cell density reached 90%, the cells were treated with ethanol or actinonin (final concentration of 10 µg/ml) for 10 h. Thereafter, the cells were harvested by trypsinization and then lysed in RIPA buffer (89900; Thermo Fisher Scientific) containing a protease inhibitor cocktail and 20 µg/ml actinonin, incubated on ice for 20 min, sonicated briefly, and centrifuged at 12,000*g* for 10 min at 4°C to obtain clear supernatants. The supernatants were diluted to 2 μg/μl, mixed with 4× SDS sample buffer in a ratio of 3:1 (v/v), and heated at 70°C for 10 min to prepare samples for immunoblotting.

### 2.4. Immunoblotting with antisera

15 μg of each sample was subjected to immunoblotting with anti-fMGSGC, anti-fMdPEG4C, or anti-fMXC sera. For human cell extracts, samples were also subjected to immunoblotting with anti-tubulin (T5168; Merck). Horse reddish peroxidase (HRP)-conjugated goat anti-rabbit IgG (BR1706515; Bio-Rad) or anti-mouse IgG (BR1706516; Bio-Rad) was used as the secondary antibody. Immunoblots were developed with ECL substrate (1705061; Bio-Rad) and visualized in chemiluminescence mode using Amersham Imager 680 (AI680; GE HealthCare, IL, USA).

### 2.5. Purification of MD-D2^3–11^-GST and fMD-D2^3–11^-GST

AG100a (*ΔarcAB*) *E. coli* cells (35) were transformed with a plasmid expressing MD-D2^3–11^-GST (MD, Met-Asp; D2^3-11^, amino acids 3-11 of the D2 protein from *Chlamydomonas reinhardtii*; GST, glutathione-S-transferase) (17,25) in pET23a(+). Expression of the indicated proteins was induced by adding isopropyl β-D-1-thiogalactopyranoside (IPTG) (I2481C; GoldBio, MO, USA) (0.5 mM final concentration) and subsequently culturing at 30°C for 4 h. To produce Nt-formylated fMD-D2^3–11^-GST, actinonin (final concentration of 2 μg/ml) was treated together with IPTG. Purification of MD-D2^3–11^-GST and fMD-D2^3–11^-GST was performed using Glutathione Sepharose resin (17075605; GE Healthcare) as described previously (25).

### 2.6. fMet Enzyme-linked immunosorbent assay (ELISA)

Each protein was prepared at concentrations of 1000 nM, 100 nM, 10 nM, 1 nM, and 0.1 nM in 50 mM carbonate coating buffer (45 mM NaHCO₃ and 5 mM Na₂CO₃). A total of 100 µl of the various concentrations of MD-D2^3-11^-GST and fMD-D2^3-11^-GST protein samples were loaded into each well of a 96-well white, non-transparent plate and coated overnight at 4°C. Afterward, each well was washed three times with 200 µl of PBS-T (137 mM NaCl, 2.7 mM KCl, 10 mM Na₂HPO₄, 1.8 mM KH₂PO₄, and 0.1% Tween 20), followed by incubation with 200 µl of blocking solution [3% bovine serum albumin (BSA) (10735086001; Merck) in PBS-T at 37 °C for 1 h. The wells were washed three more times with PBS-T. Unpurified anti-fMGSGC, anti-fMdPEG4C, and anti-fMXC sera were diluted 1:1,000 in the blocking solution and incubated in each well at 37°C for 2 h. After washing three times with PBS-T, the wells were treated with a secondary antibody (goat anti-rabbit IgG HRP conjugate, diluted 1:10,000 in blocking solution) at 37°C for 1 h. Following five washes with PBS-T, luminescence detection was performed using the SuperSignal ELISA Femto Substrate reagent (37075; Thermo Fisher Scientific) and a Promega GloMax microplate reader (Promega, WI, USA).

## 3. Results & discussion

### 3.1. Generation of antisera against fMet

In this study, we aimed to raise pan-fMet-specific antibodies to develop a more sensitive tool for detecting fMet-peptides or proteins. We designed, synthesized, and utilized three distinct fMet-bearing antigen peptides: fMGSGC, fMdPEG4C, and fMXC for antibody generation (Figure 1A). Each antigen peptide features Nt-fMet as the epitope and C-terminal Cys as a conjugatable residue, differing only in their linkers between fMet and Cys. The linker for the first antigen peptide, referenced in a previous study (2), is Gly-Ser-Gly, which provides flexibility and low immunogenicity (36). The second antigen peptide employs a dPEG4 (four repeats of discrete polyethylene glycol) chemical linker with high flexibility and stability (37). The final antigen peptide incorporates one of 20 amino acids at the linker position. To account for the sequence diversity of natural fMet-peptides and fMet-proteins in biological systems (22) (25) (2), we designed a mixture of antigen peptides that include all 20 amino acids as the second amino acid. Each antigen peptide was conjugated to the KLH carrier protein and used to immunize rabbits, ultimately yielding polyclonal antisera containing fMet-specific antibodies (Figure 1B).

**Figure 1.**
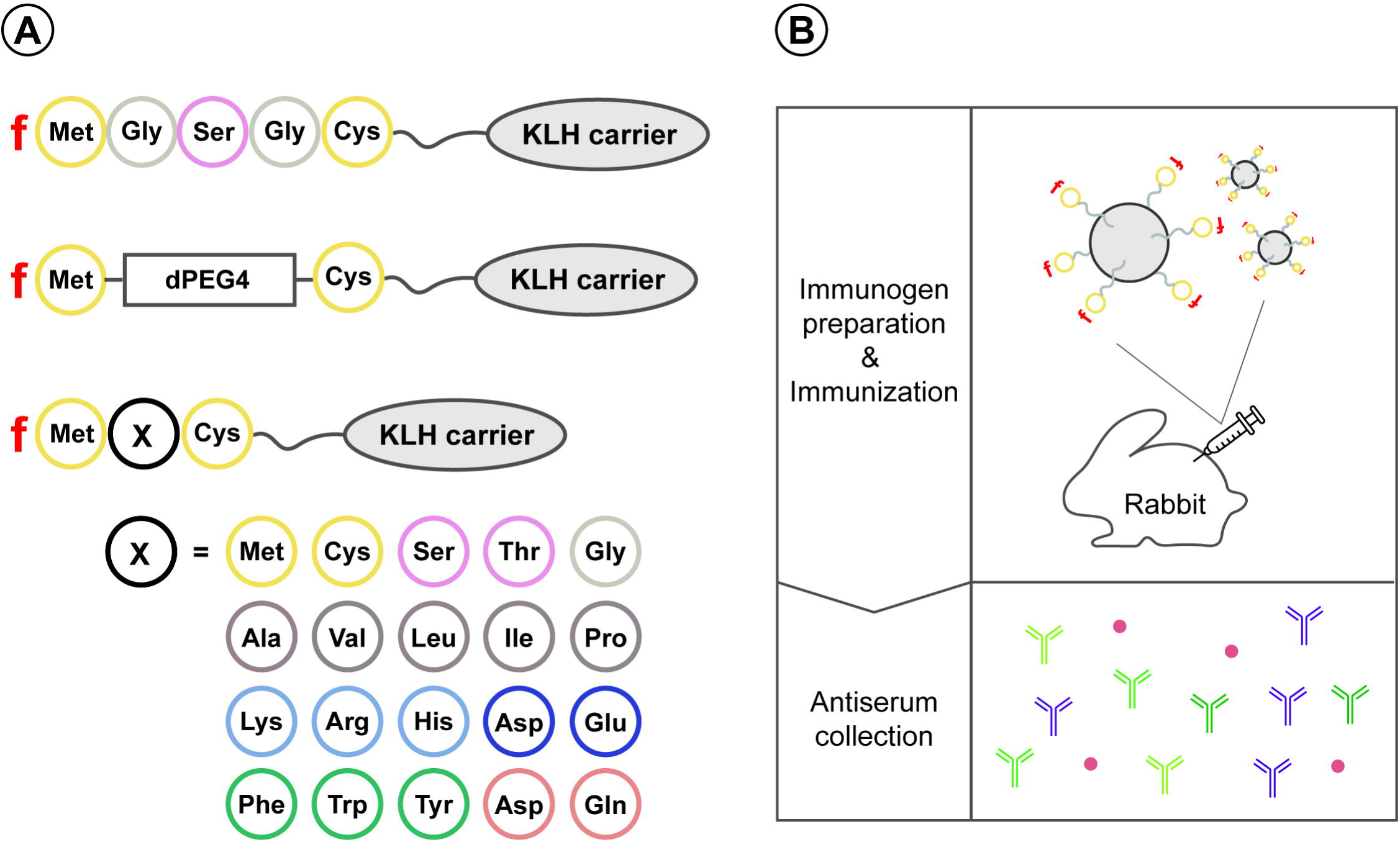
Generation of rabbit polyclonal antisera against Nt-fMet. **(A)** Synthetic fMet-peptide antigens. All antigens were designed to contain Nt-fMet and a conjugatable Cys residue, and they were conjugated to a KLH carrier protein. Each antigen features a distinct linker between fMet and Cys, including a GSG flexible linker, a dPEG4 chemical linker, or one of 20 amino acids. **(B)** Schematic of antisera generation. The three antigens were separately immunized to rabbits to produce distinct antisera for each antigen.

### 3.2 Detection of native fMet-proteins in E. coli

To evaluate the broad specificity and sensitivity of the antisera, we initially used *E. coli* cell extracts. Since nearly all proteins in bacteria are synthesized with Nt-fMet and subsequently deformylated by PDF deformylase, *E. coli* normally maintains low levels of fMet-proteins (20,22). However, treatment with the PDF inhibitor actinonin dramatically increases the levels of fMet-proteins (22). Therefore, we cultured *E. coli* with or without actinonin treatment to obtain *E. coli* extracts with high or low amounts of fMet-proteins and performed immunoblotting with these extracts and the generated antisera (Figure 2). While CBB staining confirmed equal amounts of loaded proteins across the samples (Figure 2A), among the antisera tested, those produced using the fMGSGC antigen and the mixed fMXC antigen displayed numerous signals of fMet-proteins upon actinonin treatment (Figure 2B & D). While CBB staining confirmed equal amounts of loaded proteins across the samples (Figure 2A), among the antisera tested, those generated using the fMGSGC antigen and the mixed fMXC antigen displayed numerous signals for fMet-proteins following actinonin treatment (Figures 2B & D). These results indicate the dramatic increase in both the species and amounts of Nt-formylated proteins. In contrast, the fMdPEG4C antisera exhibited relatively low binding capacity for fMet-proteins compared with the fMGSGC and fMXC antisera (Figure 2C), suggesting that the fMdPEG4C antigen is unsuitable for generating an effective anti-fMet-specific antibody.

**Figure 2.**
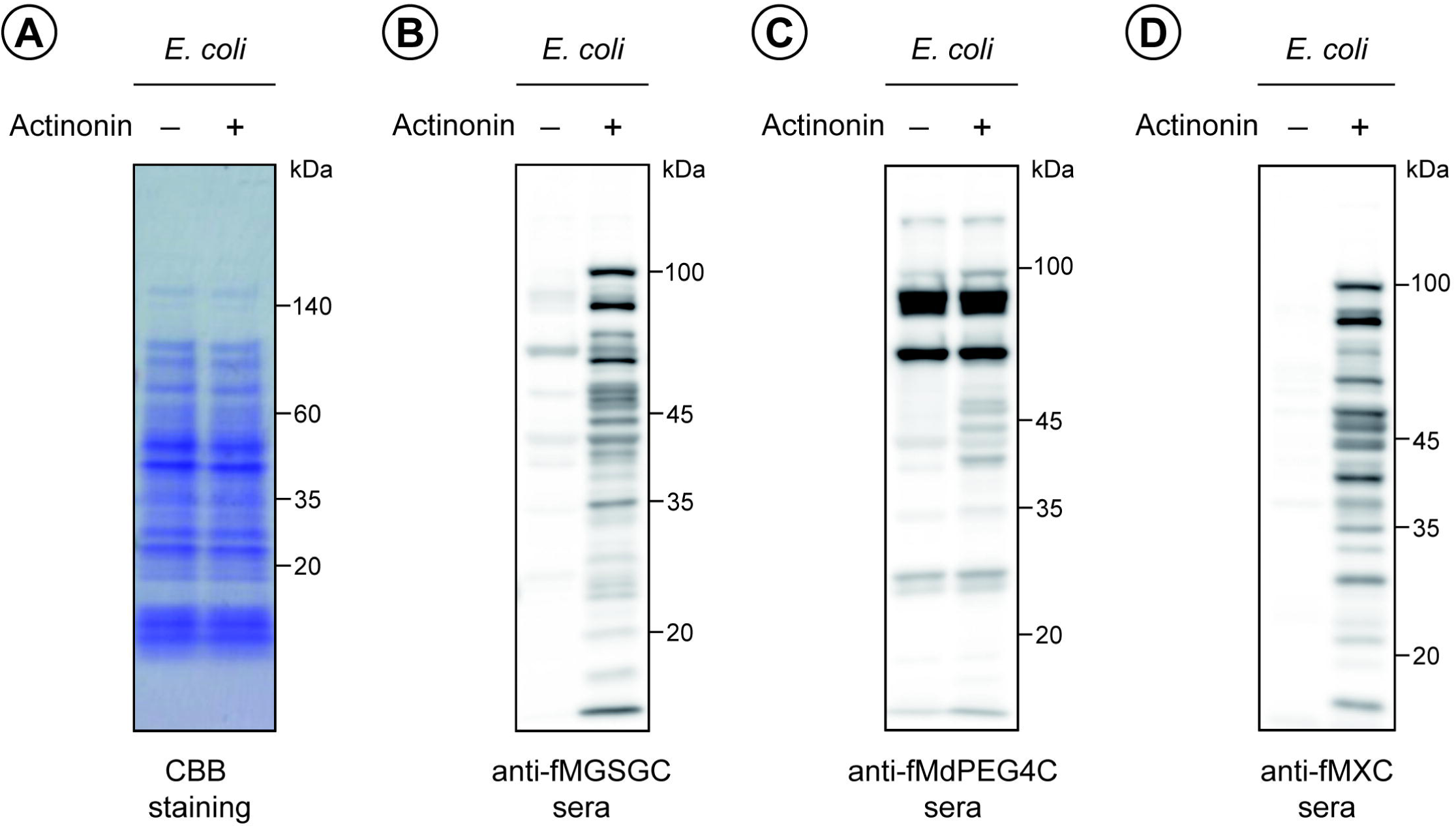
Detection of native fMet-proteins in *E. coli.* **(A)** Coomassie brilliant blue (CBB) staining of *E. coli* cell extracts, treated with or without the PDF inhibitor, actinonin (2 µg/ml) for 30 min, showing equal protein loading. **(B)** Immunoblotting of the same extracts from (A) using anti-fMGSGC sera. **(C)** Same as in (B), but with anti-fMdPEG4C sera. **(D)** Same as in (B), but with anti-fMXC sera.

### 3.2 Detection of native fMet-proteins in human HK2 cells

Next, we analyzed the expression of fMet-proteins in human cells using the antisera. Among human proteins, only 13 mitochondrial DNA-encoded proteins mainly retain Nt-fMet due to mitochondrial protein synthesis involving MTFMT and mitochondrial fMet-tRNAi (15) (27). These mitochondrial DNA-encoded proteins are also deformylated by mitochondrial peptide deformylase (mtPDF) (38), limiting the species and levels of fMet-proteins within the mitochondria. Consequently, human cells are expected to exhibit a milder response to actinonin treatment compared with bacterial cells regarding the number and kinds of fMet-proteins, despite their deformylation.

To effectively evaluate the raised antisera against fMet-proteins in human cells, we employed HK2 cells, which have a relatively high mitochondrial content compared to many other cell types (39) (40). These cells, derived from human kidney tissue, were cultured in the absence or presence of actinonin and subjected to immunoblotting with the three different antisera (Figure 3). The antisera against fMGSGC and fMdPEG4C did not show a significant increase in fMet-protein signals following actinonin treatment; however, the fMGSGC antisera did mildly detect a few fMet-proteins in the actinonin-treated samples with 20–50 kDa molecular masses (Figure 3B & C). In contrast, the fMXC antisera effectively identified fMet-proteins in HK2 cells after actinonin treatment (Figure 3D). Overall, the fMXC antisera demonstrated enhanced sensitivity to fMet-proteins in immunoblotting with the mitochondria-rich HK2 cells.

**Figure 3.**
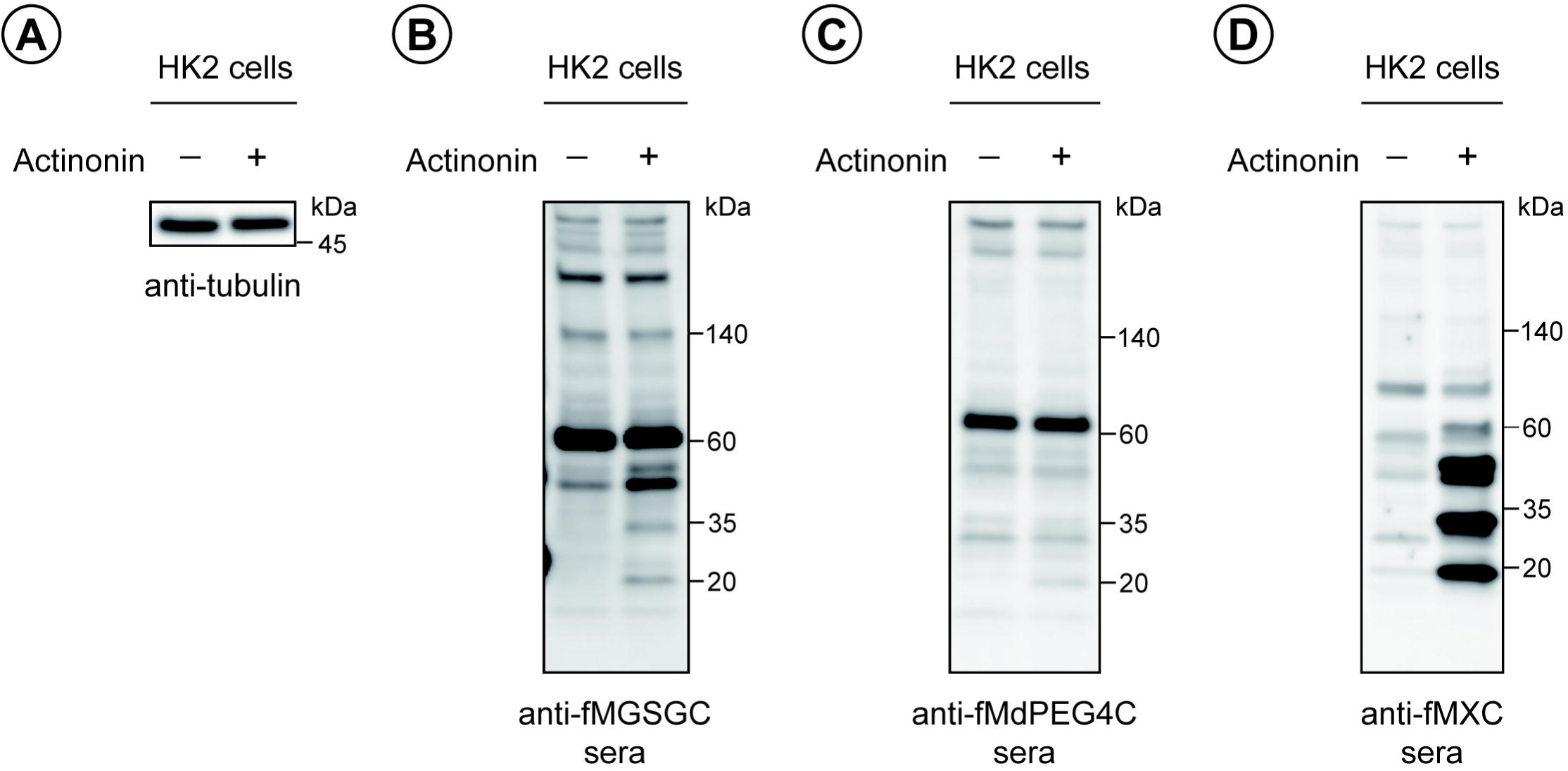
Detection of native fMet-proteins in human cells. **(A)** Immunoblotting of human HK2 cell extracts using anti-tubulin. HK2 cells were cultured in the absence or presence of actinonin (10 µg/ml) for 10 h. Tubulin serves as a loading control to indicate equal protein amounts. **(B)** Immunoblotting of the same extracts as in (A), but using anti-fMGSGC sera. **(C)** Same as in (B), but with anti-fMdPEG4C sera. **(D)** Same as in (B), but with anti-fMXC sera.

### 3.2 Quantification of fMet-proteins using ELISA

To evaluate the performance of the antisera in ELISA, previously characterized Nt-formylated fMD-D2^3-11^-GST and its unformylated counterpart MD-D2^3-11^-GST were coated onto a plate, and individual antisera were applied. All antisera distinguished fMD-D2^3-11^-GST and MD-D2^3-11^-GST with varying degrees (Figure 4). Among them, the fMXC antisera produced the highest luminescence signal with fMD-D2^3-11^-GST, indicating strong reactivity. The fMGSGC antisera generated a lower luminescence signal than fMXC, but still higher than fMdPEG4C which barely produced a fMet-specific signal. Notably, the fMXC antisera represented superior sensitivity, particularly at low antigen concentrations (10–1 nM) (Figure 4). Given that both the immunoblotting and ELISA assays were performed using crude antisera without further purification or concentration, the fMXC antisera appear to contain a significant amount of pan-fMet-specific antibodies.

**Figure 4.**
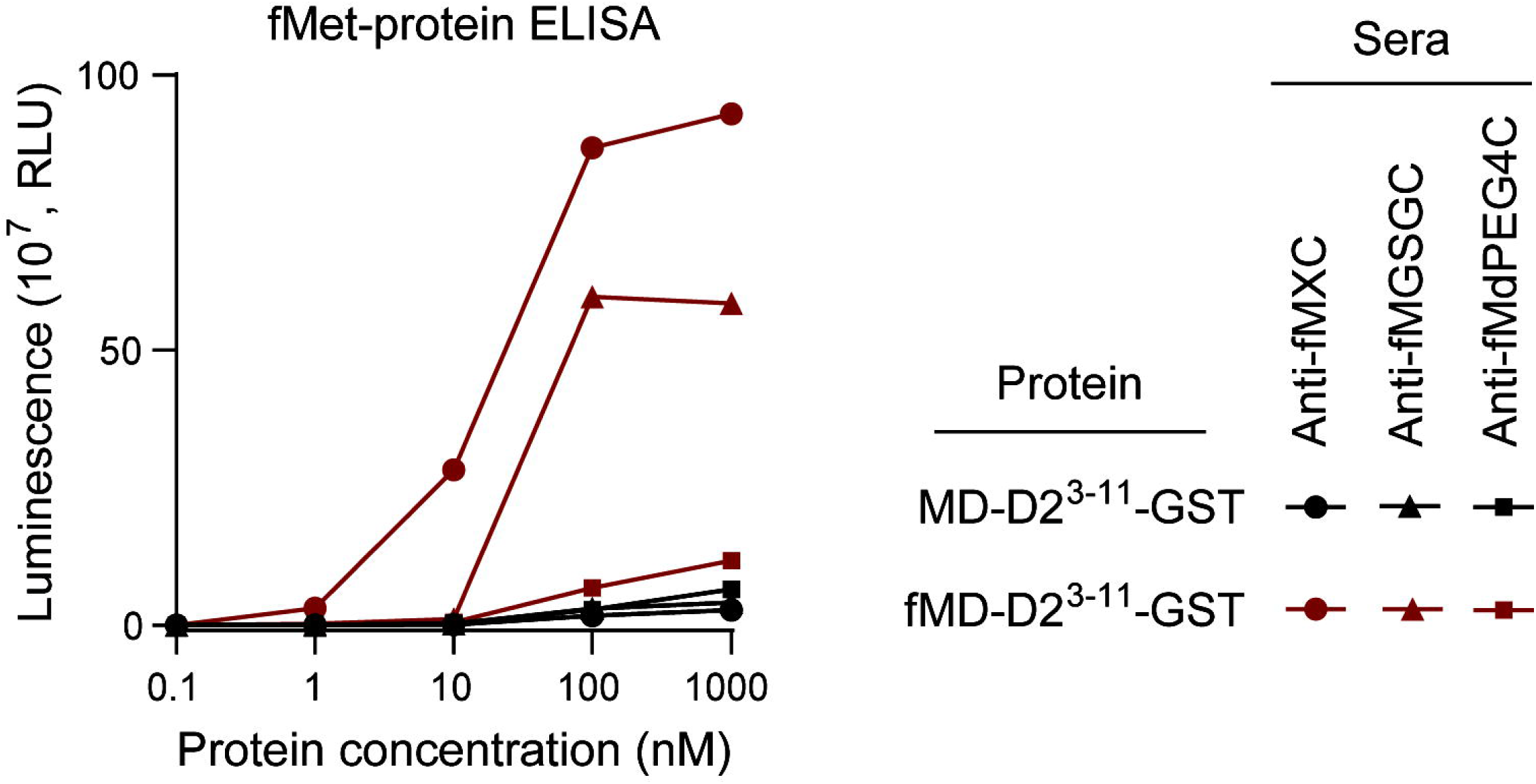
Evaluation of anti-fMet antisera using ELISA. Individual antisera were tested against Nt- formylated fMD-D2^3-11^-GST and its unformylated counterpart, MD-D2^3-11^-GST, which was serially diluted and coated onto plates. The binding affinity of the antisera was measured by applying an HRP-conjugated anti-rabbit secondary antibody, followed by luminescence detection using a luminometer. Results were expressed in relative light unit (RLU).

## 4. Conclusion

Three different fMet-bearing antigen peptides (fMGSGC, fMdPEG4C, and fMXC) were utilized for the development of pan-fMet-specific antibodies. High levels of fMet-proteins from *E. coli* were sensitively detected by both the anti-fMGSGC and anti-fMXC sera, but were barely detected by the anti-fMdPEG4C sera. However, when detecting the much lower expressed fMet-proteins from mitochondria-rich HK2 human cells, the anti-fMXC serum outperformed both the anti-fMGSGC and anti-fMdPEG4C sera. Consistently, the fMXC antisera demonstrated the highest sensitivity in the ELISA assay against fMet-proteins. In conclusion, the newly designed fMXC mixed peptide antigen significantly enhanced the production of an anti-pan-fMet-specific antibody. Given the wide variety of amino acid sequences in naturally occurring fMet-peptides or fMet-proteins, the use of a mixed peptide antigen proved superior for generating an anti-pan-fMet-specific antibody, a feat not achieved with single peptide antigens.

## 5. Future perspective

The anti-pan-fMet-specific antibody developed using the fMXC mixed peptide antigen will provide a more sensitive, convenient, and cost-effective tool for detecting fMet-peptides and fMet-proteins across various organisms, from bacteria to humans, as well as in both physiological and pathological conditions. To ensure a sustainable supply, monoclonal antibody production will be necessary and is planned for future efforts. Furthermore, the design strategy for the mixed peptide antigen described in this study can be extended to develop other anti-pan-specific antibodies targeting Nt-modifications through acetylation, methylation, oxidation, arginylation, phosphorylation, lipidation, etc.

## Acknowledgments

We thank our present and former members for their advice and assistance.

## Author contributions

D.K., K.-S.P., and C.-S.H. designed the study. D.K. performed the experiments and analyzed the data.

D.K. and C.-S.H. wrote the manuscript. C.-S.H. and K.-S.P. acquired funding. All authors participated in the discussions of the results and agreed with the content of the manuscript.

## Financial disclosure

This study was supported by Korean government (MSIP) Grants, NRF-2020R1A3B2078127 (to C.-S.H.) and NRF-RS-2024-00397929 (to K.-S.P), and Korea University grant, K2403431 (to C.-S.H.).

## Competing interests disclosure

All authors have no competing interests with the contents of this article.

## Writing disclosure

No writing assistance was utilized in the production of this manuscript, except for ChatGPT, which was used for English language editing.

## Ethical conduct of research

The are no applicable issues for this study.

## Access to anti-fMet antisera

The anti-fMet sera are currently in the pre-commercialization. Please contact the corresponding author C.-S.H for any information regarding access to the anti-fMet antisera.

## Article highlights

### Background

- Formylmethionine (fMet) acts as a key initiator of protein synthesis in both bacteria and eukaryotes, but detecting fMet-bearing polypeptides has been difficult due to the absence of effective antibodies.
- This study aims to develop a more sensitive anti-pan-fMet-specific antibody to detect fMet-bearing proteins and their derivatives across different species.

### Materials and methods

- Three distinct fMet-bearing peptide antigens—fMGSGC, fMdPEG4C, and fMXC (X, any amino acids)—were conjugated to KLH and used to generate rabbit polyclonal antisera.
- These antibodies were tested for sensitivity and specificity via immunoblotting of *E. coli* and human H2K cell lysates, as well as ELISA tests on Nt-formylated and unformylated proteins.

### Results & discussion

- Antisera against the fMXC antigen showed the highest sensitivity and specificity for detecting fMet-bearing proteins, outperforming those raised agains fMGSGC and fMdPEG4C.
- The fMXC-based antibody efficiently detected fMet-bearing proteins in both bacterial and human cell extracts, demonstrating exceptional sensitivity in ELISA for fMet-bearing proteins.

### Conclusion

- The newly developed anti-pan-fMet-specific antibody using the fMXC antigen provides a sensitive and cost-effective tool for detecting fMet-bearing proteins and their derivatives.
- This advancement opens avenues for broader applications in studying the synthesis of fMet-bearing polypeptides and cellular metabolic fates of the fMet-bearing polypeptides across species, from bacteria to humans.
- The mixed peptide antigen strategy demonstrated here would serve as a model for the future development of antibodies to other Nt-modifications.

## Notes

### Competing Interest Statement

The authors have declared no competing interest.

